# Neutrophils contribute to ER stress in lung epithelial cells in the pristane-induced diffuse alveolar hemorrhage mouse model

**DOI:** 10.1101/2021.10.01.462788

**Authors:** Gantsetseg Tumurkhuu, Duygu Ercan Laguna, Richard E. Moore, Jorge Contreras, Gabriela De Los Santos, Luisa Akaveka, Erica N. Montano, Yizhou Wang, Mariko Ishimori, Swamy Venuturupalli, Lindsy Forbess, Barry R. Stripp, Daniel J. Wallace, Caroline A. Jefferies

## Abstract

Diffuse alveolar hemorrhage (DAH), although rare, is a life-threatening complication of systemic lupus erythematosus (SLE). Little is known about the pathophysiology of DAH in humans, although increasingly neutrophils, NETosis and inflammatory monocytes have been shown to play an important role in the pristane-induced model of SLE which develops lung hemorrhage and recapitulates many of the pathologic features of human DAH. Using this experimental model, we asked whether endoplasmic reticulum (ER) stress played a role in driving the pathology of pulmonary hemorrhage and what role infiltrating neutrophils had in this process. Analysis of lung tissue from pristane-treated mice showed genes associated with ER stress and NETosis were increased in a time-dependent manner and reflected the timing of CD11b^+^Ly6G^+^ neutrophil accumulation in the lung. Using precision cut lung slices from untreated mice we observed that neutrophils isolated from the peritoneal cavity of pristane-treated mice could directly induce the expression of genes associated with ER stress, namely *Chop* and *Bip*. Mice which had myeloid-specific deletion of PAD4 were generated and treated with pristane to assess the involvement of PAD4 and PAD4-dependent NET formation in pristane-induced lung inflammation. Specific deletion of PAD4 in myeloid cells resulted in decreased expression of ER stress genes in the pristane model, with accompanying reduction in IFN-driven genes and pathology. Lastly, coculture experiments of human neutrophils and human lung epithelial cell line (BEAS-2b) showed neutrophils from SLE patients induced significantly more ER stress and interferon-stimulated genes in epithelial cells compared to healthy control neutrophils. These results support a pathogenic role of neutrophils and NETs in lung injury during pristane-induced DAH through the induction of ER stress response and suggest that overactivation of neutrophils in SLE and NETosis may underlie development of DAH.

## Introduction

Diffuse alveolar hemorrhage (DAH) is a rare but severely life-threatening complication observed in patients with systemic lupus erythematosus (SLE) and other autoimmune diseases including anti-neutrophil cytoplasmic antibody (ANCA) vasculitis (1, 2). DAH is driven by injury to the alveolar capillary bed basement membrane leading to accumulation of red blood cells in the alveolar space. What causes injury to the basement membrane is specific to the systemic disease that underlies DAH. Specifically in SLE- and ANCA-associated vasculitis, immune complex deposition, complement activation, and neutrophil infiltration of the alveolar wall are the key histopathologic findings (3).

Type I IFN-producing neutrophils have come to the forefront in driving both inflammation and exacerbating tissue pathology in lupus (4-7). Termed low density neutrophils (LDNs), they respond to stimuli such as immune complexes with a much higher propensity to release neutrophil extracellular traps (NETs), a process called NETosis, compared to normal density neutrophils (8, 9). NETosis is a critical physiological response to bacterial and fungal infections, aiding to trap, immobilize, and help prevent the spread of infection (10). NETs are also triggered by cytokines, immune complexes and autoantibodies (11). The enzyme peptidylarginine deiminase 4 (PAD4) is expressed in mouse and human neutrophils and catalyzes the citrullination of histones leading to chromatin decondensation, a hallmark of NETosis. PAD4 has been proposed as a critical enzyme in the formation of NETs and its deletion or inhibition leads to an abrogation of NETosis in various contexts including bacterial infection, atherosclerosis, and the MRL/lpr SLE model (11-14). In addition, myeloid-specific deletion of PAD4 results in an inability of neutrophils from these mice to induce NETosis in response to the calcium ionophore A23187 and a reduction in inflammation and atherosclerosis in the context of ApoE-deficiency (15). The release of DNA and nuclear histones during NETosis drives inflammation and type I IFN production through activation of RNA/DNA sensing pathways (4). Although these mechanisms are vital for anti-microbial defenses, inappropriate activation exacerbates SLE pathology and acute lung injury (16). Recent evidence using pristane-induced lupus model where the mice develop DAH demonstrates that NETosis is an important contributor to lung injury (11). In this model, neutrophils accumulate in the lungs of mice following a single intraperitoneal injection of the hydrocarbon oil pristane. Enhanced NETosis is observed in the lungs of pristane-treated mice and treatment with recombinant DNase I reduced NET formation in the lung with a subsequent reduction in lung injury in C57BL/6 mice (17). Thus, NETs may play key roles in driving pristane-induced lung injury (17).

Additional pathways have also been shown to contribute to acute and chronic lung injury. For example, evidence from the monogenic autoimmune disorder COPA syndrome, which presents frequently as pulmonary hemorrhage, suggests that endoplasmic reticulum (ER) stress may be a key driver of lung injury in autoimmune disease (18). ER stress is sensed by ER-resident transmembrane proteins such as IRE1, ATF6 and PERK. For example, unfolded proteins that initiate ER stress bind to BIP which then dissociates from IRE1 and PERK to promote their phosphorylation and activation. IRE1 and PERK activation triggers cell protective mechanisms such as the synthesis of the ER chaperone protein CHOP via the generation of spliced *Xbp1* transcripts, downstream of IRE1 and/or ATF6 activation. Increased expression of *Bip* or *Chop* and higher spliced to unspliced *Xbp1* transcript ratios are therefore useful markers of enhanced ER stress. Recent evidence suggests that activation of the ER stress response can act as a danger signal to augment the inflammatory responses in bronchial epithelial cells (19). Importantly, inhibiting ER stress can alleviate LPS-induced lung injury in mice (20), suggesting the importance of this pathway in inflammatory lung disease. Recently, crosstalk between NETs and ER stress has been observed in a model of sepsis-induced intestinal injury, with inhibition of NETosis and ER stress reducing intestinal epithelial cell monolayer barrier disruption, suggesting that neutrophils and NETosis contribute to ER stress in intestinal epithelial cells (21).

We hypothesized that NETs and NETosis directly drive ER stress in lung epithelial cells and contribute to acute lung injury in the pristane mouse model. In keeping with this we observed that expression of ER stress genes (*Bip* and *Chop*) increased in a time-dependent manner and closely matched the influx of neutrophils into the lungs of pristane-treated mice. To assess the importance of NETosis in driving lung inflammation in this model, we generated mice with myeloid specific deletion of PAD4 (LysMCrePAD4^fl/fl^ mice) and found that these mice showed reduced lung pathology and expression of ER stress and ISGs when treated with pristane compared to C57BL/6 (WT) mice. Co-culturing precision-cut lung slices from WT mice with neutrophils from pristane-treated PAD4^fl/fl^ mice drove expression of both ER stress markers and ISGs whereas (11-14) neutrophils from LysMCrePAD4^fl/fl^ mice failed to do so, supporting NETosis as a mechanism for driving ER stress. Because low-density neutrophils from SLE patients undergo spontaneous NETosis *ex vivo* (9), we hypothesized that neutrophils from SLE patients would drive ER stress in alveolar epithelial cells when compared with neutrophils from healthy controls (HC). As expected, SLE neutrophils, but not HC neutrophils, cocultured with an alveolar epithelial cell line induced high levels of ER stress and genes associated with inflammation in the lung such as *Il8* and *Il16*, suggesting an important role for neutrophils in driving ER stress and inflammation in lung epithelial cells in the context of SLE. Taken together our results demonstrate the importance of neutrophils and NETs during lung injury in the pristane mouse model and in their implications in human SLE-related DAH.

## MATERIALS and METHODS

### Patient Samples

All SLE patients (as per ACR diagnostic criteria) were recruited from Cedars-Sinai Medical Centre, CA, USA. The Systemic Lupus Erythematosus Disease Activity Index (SLEDAI) score was determined for each patient at the time of the blood draw. Age- and sex-matched healthy donors who had no history of autoimmune diseases or treatment with immunosuppressive agents were included. All participants provided informed written consent and the study received prior approval from the relevant institutional ethics review boards (IRB protocol 19627).

### Isolation of PBMCs and Cellular Subsets

Peripheral blood mononuclear cells (PBMCs) were separated from whole blood by density-gradient centrifugation with Ficoll-Paque Plus (GE Healthcare). Neutrophils were isolated using the EasySep Direct Human Neutrophil Isolation Kit (STEMCELL Technologies, 19666) according to manufacturer’s protocol.

### Cell Culture

Primary BEAS-2B (ATCC® CRL-9609TM) (normal human airway epithelial cells) as well as all the basal media and growth supplements were obtained from Lonza (Walkersville, MD). Cells were cultivated according to the instructions of the manufacturer on plastic dishes or flasks (BD Bioscience, Heidelberg, Germany). Passage number was kept to less than four passages from original stocks.

### Gene Expression Analysis

Total RNA was extracted using TRIzol reagent (Invitrogen, Carlsbad, CA), and cDNA was synthesized using the iScript Reverse Transcription Supermix Kit (Bio-Rad) according to the manufacturer’s recommendations with the PerfecTa SYBR Green PCR Kit (Quanta, 95072-012) as per the manufacturer’s recommendations. Real-time PCR data were analyzed using the 2^-ΔΔCt^ method and gene expression was normalized to *GAPDH* or *18s*. Primaer details given in supplemental table 1.

### Animal Studies

All animal experiments were performed according to the guidelines and approved protocols of the Cedars-Sinai Medical Center Institutional Animal Care and Use Committee (IACUC protocol # 6288). Cedars-Sinai Medical Center is fully accredited by the Association for Assessment and Accreditation of Laboratory Animal Care (AAALAC International) and abides by all applicable laws governing the use of laboratory animals. Laboratory animals are maintained in accordance with the applicable portions of the Animal Welfare Act and the guidelines prescribed in the DHHS publication, Guide for the Care, and Use of Laboratory Animals.

### Mice

Wild-type C57BL/6, PAD4^fl/fl^ and LysMCre-PAD4^fl/fl^ mice (C57BL/6 background), female, 6–8 weeks old, received a single intraperitoneal injection of 0.5 ml of phosphate-buffered saline (PBS) or pristane (2,6,10,14-Tetramethylpentadecane (TMPD), Sigma, P1403). Mice were sacrificed at 1 week after PBS or pristane injection.

### Flow Cytometry

The following conjugated anti-mouse antibodies were used: anti-Ly-6G (1A8), anti-CD11b (M1/70), anti-Ly-6C (ER-MP20), anti-F4/80 (BM8), anti-CD4 (GK1.5), anti-CD8a (53–6.7), anti-CD11c (3.9), and anti-B220 (RA3-6B2) (eBiosciences). Cells were incubated in CD16/32 (Fc block; BD Biosciences) prior to staining. Cell fluorescence was acquired on LSR II (BD Biosciences) and analyzed with the FlowJo software (Treestar).

### Lung Histology

The same right lower lung lobes from mice in different groups were preserved in 10% formalin for 24 hours. Then the lung tissue was embedded in paraffin wax, sliced, and stained with hematoxylin-eosin. The lung injury was observed using an Aperio ScanScope AT Turbo and image analysis was performed by ImageJ software.

### Immunohistochemistry

The mice were sacrificed and lungs were perfused with ice-cold PBS prior to removal. Lungs were fixed for 24 hours in 10% neutral buffered formalin then stored in 70% ethanol until they were embedded in paraffin and sectioned at 5μm. The slices were deparaffinized in xylene and rehydrated to graded changes of ethanol. Antigen retrieval was performed using a pressure cooker by heating the sections for 45 minutes at 95°C in sodium citrate buffer, pH 6.0. Briefly, blocking and permeabilizing reagents were added for 30 minutes followed by incubation overnight at 4°C with primary antibodies against CHOP (Thermo Fisher, MA1-250) and E-Cadherin (Cell Signaling, 24E10). Slides were then incubated in anti-mouse-AF488 (Abcam, ab150113) or anti-rabbit-AF555 (Cell Signaling Technology, 4413S) secondary antibody, respectively, and DAPI for 30 minutes. Images were captured using a Keyence BZ-X710 microscope.

### PCLS preparation and culture

Wild-type B6 mice were sacrificed by isoflurane inhalation, and lungs were perfused with 15 ml PBS at room temperature through the right ventricle. The trachea was cannulated with a 20-gauge catheter (20G × 1.00 in. BD Insyte Autoguard), and then lung lobes were inflated with 3% (w/v) 45°C prewarmed low-melting agarose (Promega). The inflated lung lobes were immediately removed and cooled on ice for 15 minutes. After lobes solidified, the left lobe was separated and sectioned into 100-μm-thick slices using a vibratome (Leica VT 1200S, Leica Microsystems). Tissue slices were then cultured in DMEM supplemented with 10% FCS, 100 μg/ml penicillin/streptomycin, and glutamine. Isolated neutrophils were applied immediately after sectioning.

### Statistics

All data are expressed as mean ± SD. Statistical differences were measured using either an Student’s unpaired t-test or 2-way analysis of variance (ANOVA) with Bonferroni post-hoc test when appropriate. Normality of data was assessed via a Shapiro–Wilk normality test. When the data analyzed was not distributed normally, we used the Mann–Whitney test or Kruskal–Wallis 1-way ANOVA with Dunn’s post-hoc test. Data analysis was performed using Prism software version 7.0a (GraphPad, San Diego, CA). A p-value of < 0.05 was considered statistically significant. Asterisks in the figures represent the following: \**p* < 0.05; \*\**p* < 0.01; \*\*\**p* < 0.001, and \*\*\*\**p* < 0.0001.

## RESULTS

### ER stress markers increased in lungs of pristane-treated mice

In the pristane model of IFN-inducible SLE, diffuse alveolar hemorrhage (DAH) develops 10-14 days post intraperitoneal injection of pristane in C57BL6 mice (22-24). Notably, the onset of DAH is preceded by an influx of neutrophils into the lungs 3-7 days post pristane injection (Figure 1A and (25)), supported by increased levels of the neutrophil-associated gene transcripts *MPO* and *ELANE* and the NETosis-associated gene, Pad4, in the lung (Supp. Figure 1). To investigate whether ER stress is involved in pristane-induced lung injury, we measured expression levels of ER stress markers at different time points in the lungs following pristane treatment. Both *Chop* and *Bip* were increased compared to untreated mice day 7 post pristane treatment (Figure 1B), mirroring the timing of neutrophil influx. Investigating which ER stress pathway might be involved at day 7 after pristane injection, we found expression of *Ire1* and the ratio of s*Xbp1* to total *Xbp1* was increased (Figure 1C), suggesting that the IRE-1/XBP-1 signaling pathway may be involved in this model. Analysis of lung sections from pristane-treated mice showed that enhanced protein levels of the ER stress marker CHOP could be observed in E-cadherin^+^ lung epithelial cells (Figure 1D).

**Figure 1.**
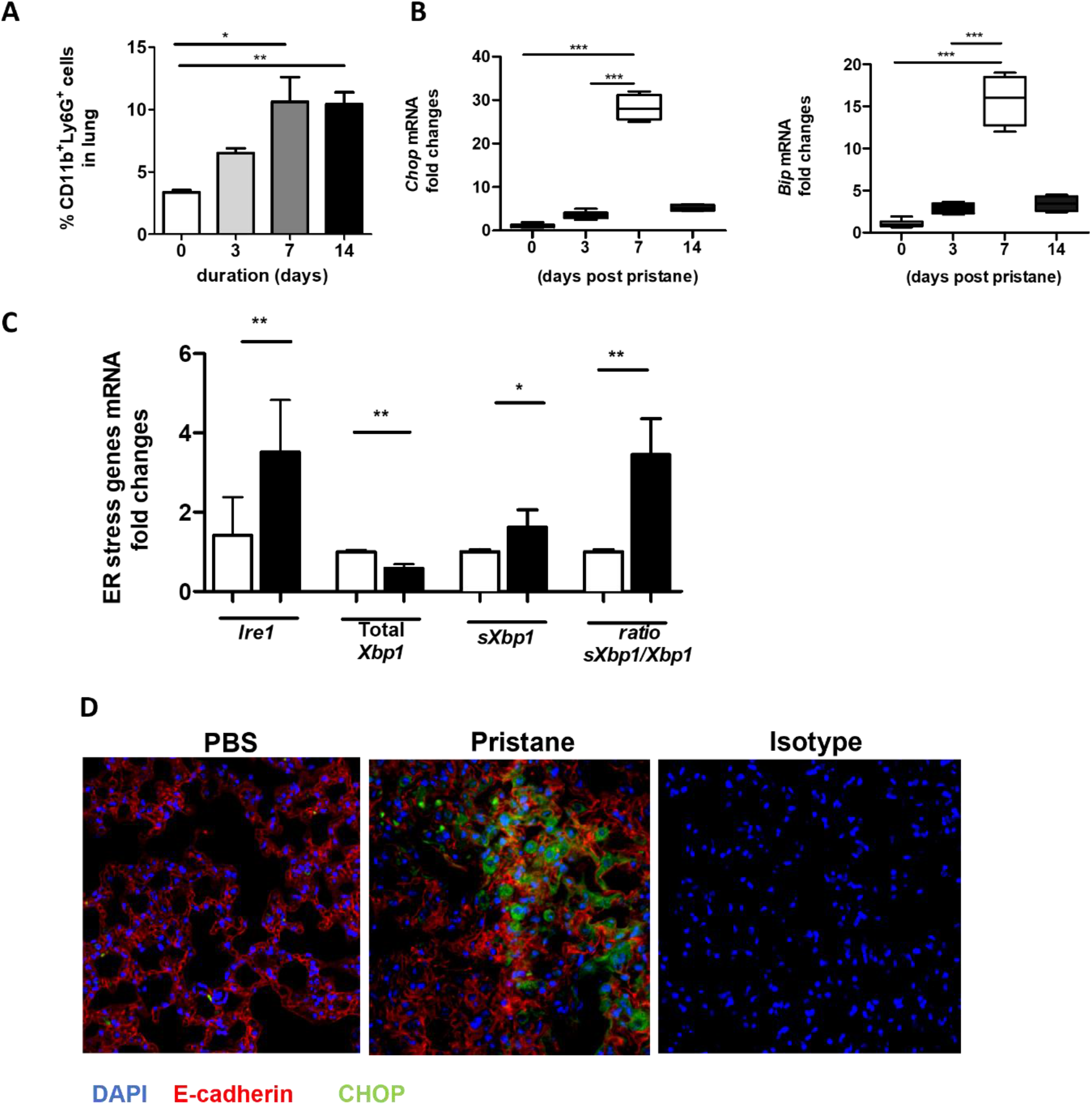
Increased levels of ER stress markers in lung of pristane mice associate with neutrophil infiltration. **(A)** Time course analysis of neutrophil infiltration in whole lung of C57BL/6 mice after pristane treatment determined by FACS. **(B)** Time course analysis of ER stress gene expression in the whole lung from indicated days after pristane treatment as measured by qPCR. **(C)** ER stress gene expression in the lung day 7 after pristane treatment as measured by qPCR. **(D)** Representative images immunostaining for CHOP (green), E-Cadherin (red), and DAPI (blue). Values are the mean ± SD. Statistical significance was determined by one-way ANOVA test. \**p* < 0.05, \*\**p* < 0.01, and \*\*\**p*, ≤ 0.001

### Pristane-activated neutrophils can drive ER stress in precision cut lung slices from untreated C57BL/6 mice

Given recent evidence suggesting that NETs drive ER stress in intestinal epithelial cells (21), we next asked if neutrophils taken from pristane-treated mice were able to drive either ER stress or inflammation when cocultured with precision cut lung slices (PCLS) from untreated C57BL/6 mice. Following pristane administration, neutrophils are recruited to the peritoneal cavity as early as 3 days post pristane treatment and remain present up to 14 days post administration (25). Neutrophils were therefore isolated from the peritoneal cavity of mice 7 days after PBS or pristane treatment and incubated with precision-cut lung slices (PCLS) generated from untreated C57BL/6 mice *ex vivo*. Compared with neutrophils from PBS-treated mice, peritoneal neutrophils from pristane-treated mice drove accumulation of ER stress markers, *Chop* and *Bip*, in addition to increased expression of *sXbp1* when co-cultured with PCLS (Figure 2A). Coculturing neutrophils from pristane-treated mice with PCLS also resulted in enhanced ISG expression (Figure 2B). Importantly, the levels of expression of both ISG and ER stress markers observed in this coculture experiment were significantly higher than pristane-treated peritoneal neutrophils cultured alone (Supp. Figure 2A). Thus, pristane-treated peritoneal neutrophils not only drive inflammation and IFN-driven responses but can also drive ER stress when cultured with PCLS.

**Figure 2.**
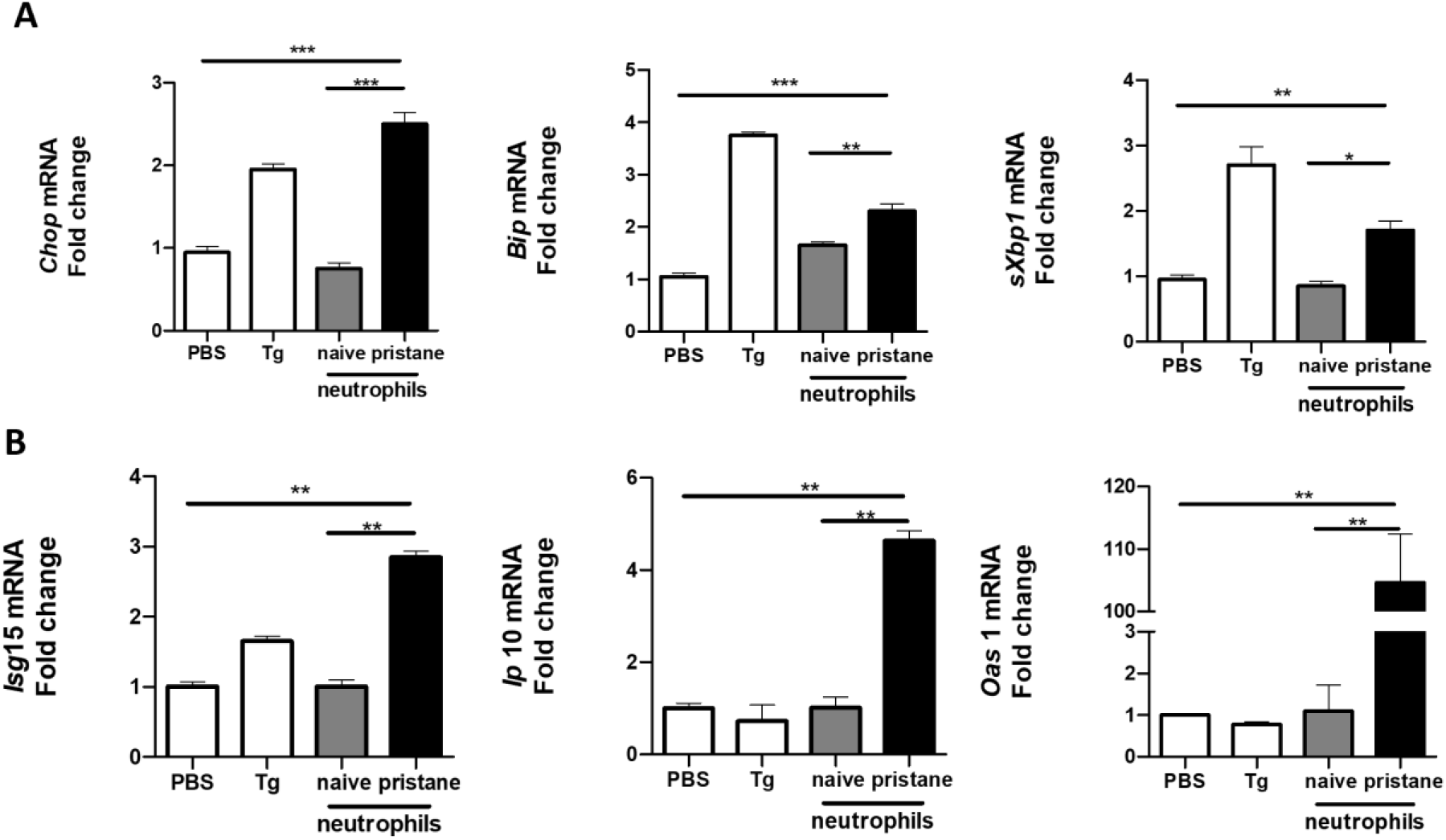
Neutrophils from pristane-treated mice Induce higher expression of ER stress and ISGs In lung cells. PCLS from untreated mice were cultured (1 slice/well) and stimulated with thapsigargin (Tg) as an ER stress positive control or cocultured with peritoneal neutrophils (4×10^6^/well) from PBS- or pristane-treated mice. Expression of **(A)** ER stress markers and **(B)** ISGs were determined from total cells by qPCR. Values are the mean ± SD. Statistical significance was determined by one-way ANOVA test. \**p* < 0.05,\*\**p* < 0.01, and \*\*\**p* ≤ 0.001

### Loss of PAD4 reduces pristane-induced lung inflammation

Histone citrullination by peptidylarginine deiminase 4 (PAD4) drives chromosome decondensation and NETosis (11). By crossing LysMCre mice to PAD4^fl/fl^ mice we asked whether blocking PAD4-dependent NETosis would reduce lung inflammation and ER stress in pristane-treated mice. Pristane-induced lung inflammation and alveolar hemorrhage was compared between C57BL6 and LysMCrePAD4^fl/fl^ mice, with significantly reduced hemorrhage observed in pristane-treated LysMCrePAD4^fl/fl^ mice compared to PAD4^fl/fl^ mice (Figure 3A), indicating that loss of *Pad4* in myeloid cells and PAD4-dependent NETosis was protective in this setting. Importantly, the ability of pristane to increase expression of ER stress markers and the ISG *Ip10* in the lung was lost in the LysMCrePAD4^fl/fl^ mice (Figure 3B). Similarly, enhanced expression of neutrophil-associated genes (*Ly6g* and *MPO*) and ISGs in peritoneal cells isolated from pristane-treated PAD4^fl/fl^ mice was abrogated in LysMCrePAD4^fl/fl^ mice (Supp. Figure 3A and B). We have previously assessed the effect of exogenous stimuli in driving lung inflammation using precision cut lung slices in an *ex vivo* culture system (25). Using this approach we next assessed whether pristane-exposed neutrophils could affect either ER stress markers or ISG expression in PCLS. Neutrophils isolated from the peritoneal cavity of pristane-treated LysMCrePAD4^fl/fl^ mice were unable to drive expression of ER stress markers or ISGs when cocultured with PCLS from untreated C57BL/6 mice (Figure 3C). Thus, PAD4-dependent NETosis is likely a primary driver of both ER stress and ISG expression in the pristane model of DAH.

**Figure 3.**
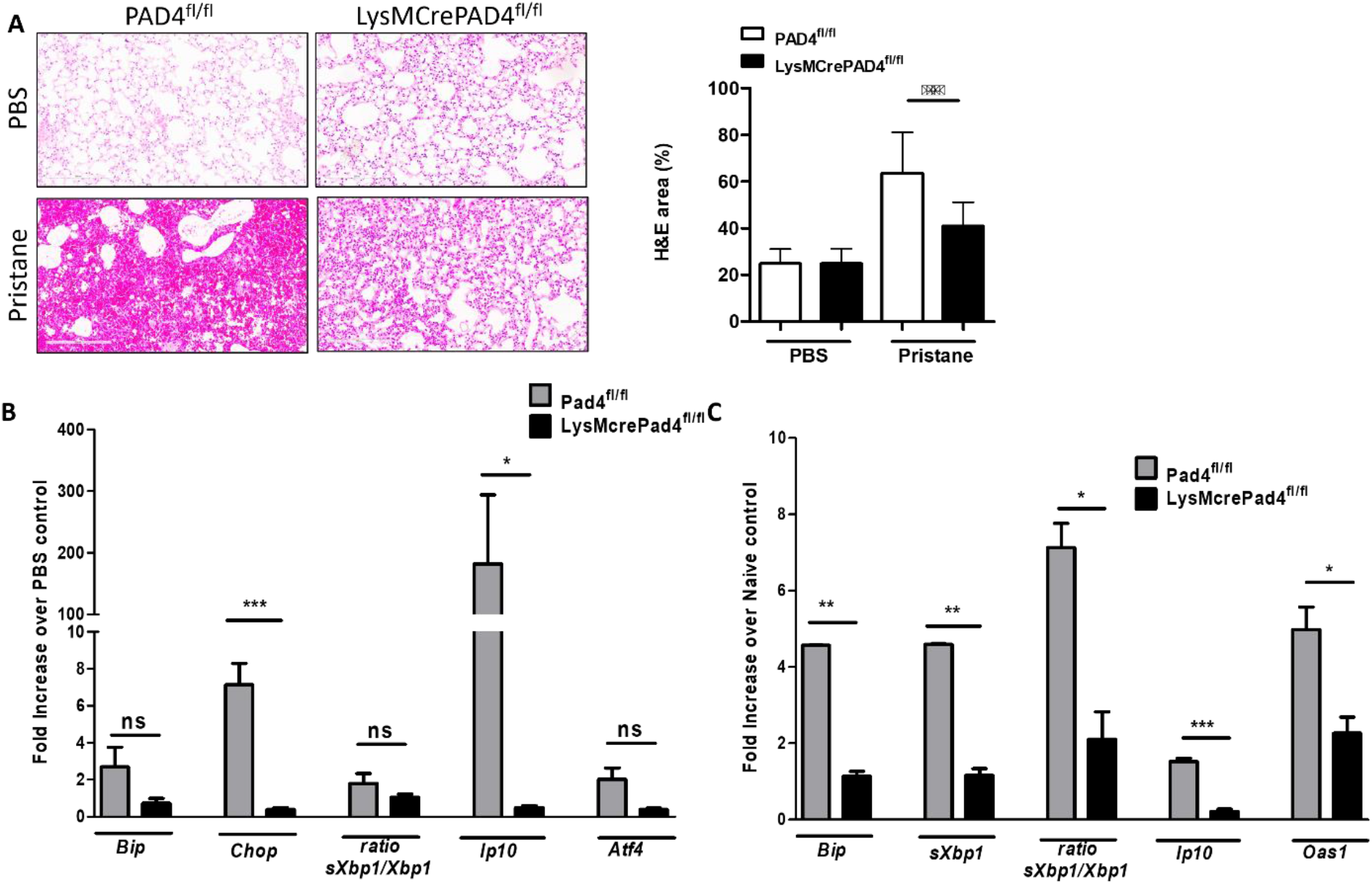
Lung inflammation and ER stress genes are driven by PAD4-expressing neutrophils 7 days post pristane treatment. **(A)** Representative haematoxylin and eosin (H&E)-stained lung sections (x200) of PBS- or pristane-treated PAD4^fl/fl^ or LysMCrePAD4^fl/fl^ mice (n=3 per group). Quantification of lung damage was determined from 10 randomly selected images. **(B)** Expression of ER stress genes and ISGs were determined by qPCR of whole lung. **(C)** PCLS were prepared and cocultured with neutrophils as described in Figure 2, and PAD4^fl/fl^ or LysMCre-PAD4^fl/fl^ peritoneal neutrophils were isolated after 7 days of PBS or pristane treatment. Values were normalized to neutrophils isolated from PBS-treated mice of their respective genotype. ER stress genes and ISGs were determined by qPCR of whole cells. Values are the mean ± SD. Statistical significance was determined by one-way ANOVA test.\**p* < 0.05, \*\**p* < 0.01, and \*\*\**p ≤* 0.001

### Neutrophils from SLE patients can drive both ISG expression and upregulate ER stress markers in lung epithelial cells

In a separate, unpublished study we have found that neutrophils from patients with moderate to active disease (SLEDAI > 4) show strong IFN signature compared to neutrophils from patients with inactive disease (Figure 4A), indicating that they may be immunostimulatory. To translate our observations in the pristane model translate to human SLE, we cocultured neutrophils isolated from active SLE patients (SLEDAI > 6) or healthy controls (HC) whole blood with BEAS-2B bronchial epithelial cells. In keeping with our observations in the pristane model, co-culture of BEAS2B cells with neutrophils from SLE patients, but not HC, resulted in increased expression of ER stress markers in the BEAS-2B cells, far above that of BEAS-2B cells or neutrophils cultured alone (Figure 4B). In addition, markers known to be associated with immune cell recruitment and lung inflammation (*IL16, IP10*, and *IL8*), were also markedly enhanced in BEAS-2B-SLE neutrophil cocultures compared with either SLE neutrophils alone or BEAS-2B-HC neutrophil cocultures (Figure 4B). Thus, neutrophils from SLE patients with active disease enhance both ER stress and inflammatory gene expression when cocultured with alveolar epithelial cells.

**Figure 4.**
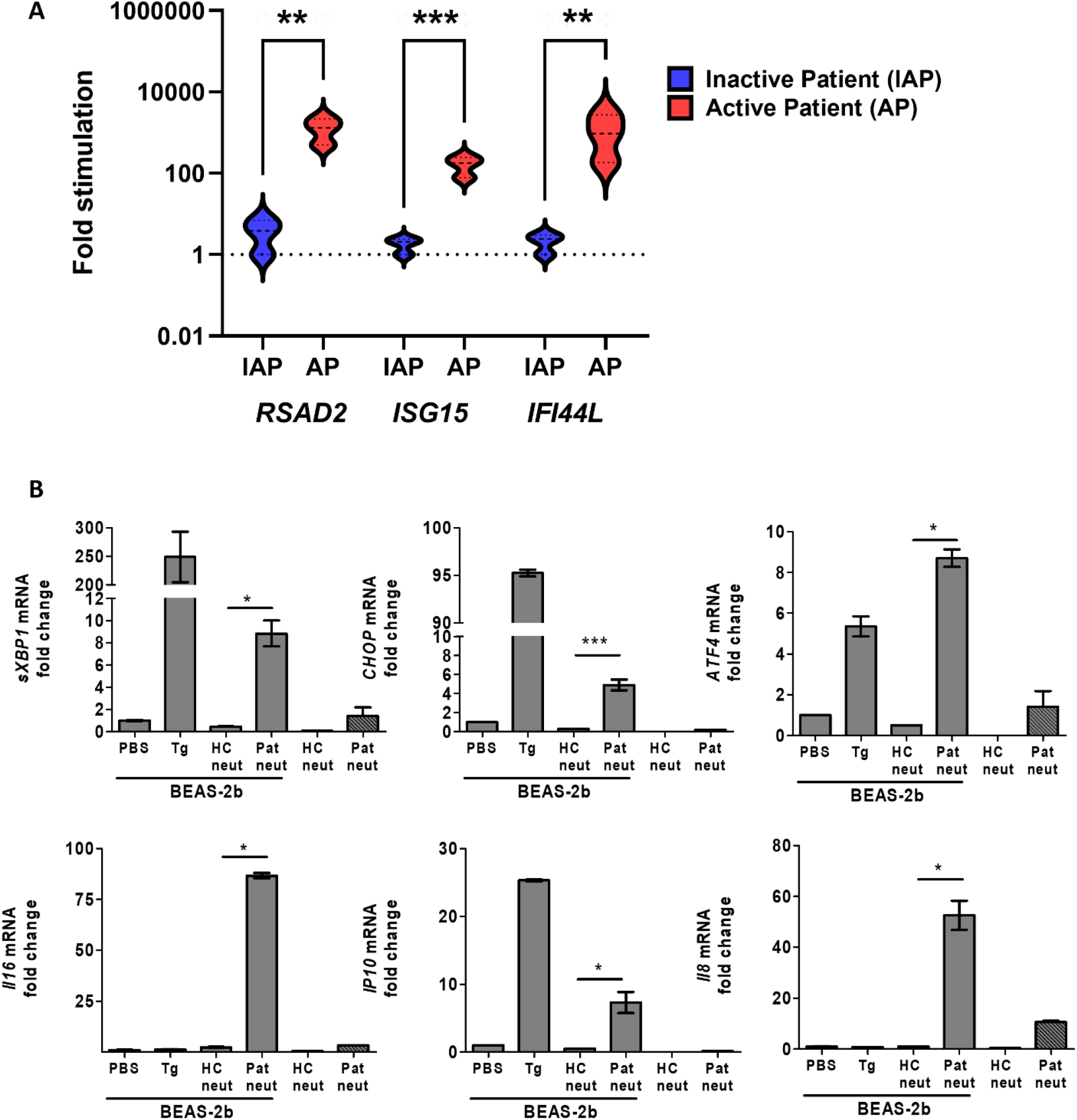
Active SLE patient neutrophils are inflammatory and induce ER stress in human BEAS-2b bronchial epithelial cells. **(A)** The expression levels of ISGs in neutrophils isolated from the whole blood of moderate to active SLE (SLEDAI > 4, AP) or inactive SLE patients (IAP) as determined by qPCR (n=3 for both groups). **(B)** Neutrophils from healthy control and active SLE patient whole blood were isolated and cultured alone or directly with BEAS-2b cells at a 1:4 ratio and incubated overnight. The expression levels of ER stress genes were determined by qPCR of whole cells. Values are the mean ± SD. Statistical significance was determined by one-way ANOVA test. \**p* < 0.05, \*\**p* < 0.01, and \*\*\**p ≤* 0.001

In summary, we have shown that neutrophils drive ER stress in the lungs of mice treated with pristane and that preventing PAD4-dependent NETosis can reverse both inflammation and induction of ER stress in pristane-treated mice. Additionally, we observed that neutrophils derived from SLE patient whole blood could directly drive ER stress in bronchial epithelial cells, suggesting ER stress as a potential pathologic mechanism in SLE.

## DISCUSSION

Increasing evidence implicates release of neutrophil extracellular traps (NETs), NETosis, may drive lung injury, specifically ER stress in epithelial cells (9, 21, 26, 27). Our study shows that not only are ER stress markers increased in the pristane model of IFN-inducible SLE, but that inhibition of NETosis using LysMCrePad4^fl/fl^ mice in this model prevented pristane-induced ER stress and, more importantly, lung inflammation. Coculture experiments demonstrated that neutrophils isolated from pristane-treated mice could directly upregulate ER stress markers in lung slices in addition to driving IFN-stimulated gene expression. Importantly, we found that neutrophils isolated from SLE patients highly upregulated inflammation markers and induced both ER stress markers and ISG expression in alveolar epithelial cells, suggesting that ER stress may be an important pathological mechanism in SLE.

Recently neutrophils and NETosis have been shown to play important roles in lung inflammation. In autoimmune lung inflammation for example, NETs have been shown to activate lung fibroblasts and promote their proliferation and differentiation into myofibroblasts and thus contribute to interstitial lung disease in patients with poly- and dermato-myositis (28, 29). In the pristane-induced DAH model, treatment of mice with recombinant DNase I reduced NET formation and reduced lung pathology (17). In keeping with this pathogenic role for NETosis in pristane-induced lung injury, we have observed that myeloid-specific deletion of *Pad4* results in reduced lung inflammation and ISG expression. Other evidence for NETosis playing a pathogenic role in lung injury models have found that inhibition of NETosis as well as disintegration of NETs by DNase I treatment was highly protective against ventilator-induced lung injury (VILI) (30). NETs were also observed in both patients and a murine model of primary graft dysfunction, with DNase I treatment again reducing lung injury (31), suggesting enhanced NETosis and the presence of NETs to play a pathologic role in lung injury. However, another study using conventional knockout mice for *Pad4* in the pristane model found that constitutive deletion of *Pad4* results in higher levels of anti-nuclear antibodies (ANAs) and inflammatory mediators (32). However, in this study mice lung inflammation was not assessed and more importantly, mice were sacrificed approximately 4 months post pristane injection, whereas DAH and lung inflammation in these mice occurs day 10-14 and resolves after approximately 28 days. Also in our short-term study, we employed myeloid specific deletion of *Pad4*, in contrast to the aforementioned study, as conventional knockout mice for *Pad4* display enhanced expression of hematopoietic multipotent progenitor numbers which may affect biology and response to agonists (33). Thus, although the potential exists that NETosis may indeed be important in helping to resolve inflammation in murine lupus models as the disease progresses, our data and data of other suggests that restricting NETosis prevents pristane-induced lung inflammation.

Type I IFNs are known to be central to the pathology of pristane-driven autoimmune disease and lung inflammation in C57BL/6 mice, with pristane-driven autoimmune disease absent in *IFNAR* knockout mice (22). Importantly, NETs have been shown to be important for induction of ISG expression in murine models of SLE and *ex vivo* analysis of SLE patient neutrophils (6, 9). The presence of dsDNA in the NETs activates DNA sensors in both immune and non-immune cells triggering IFN induction and subsequent ISG expression. Our results show that prevention of NETosis using LysMCrePad4^fl/fl^ mice results in reduction of IFN-stimulated gene expression in the lungs of pristane-treated mice. NETs have also been shown to trigger ER stress in gut epithelial cells in a model of septic shock (14). In this study, ER stress activation by NETs in intestinal epithelial cells was mediated by TLR9, suggesting that DNA in the NETs was directly triggering ER stress in these cells. Our study shows that neutrophils can directly induce ER stress in epithelial cells, with *Pad4*-deleted neutrophils unable to drive ER stress in coculture experiments. Interestingly, in the aforementioned study, TLR9-driven ER stress increased gut barrier permeability, potentially suggesting a mechanism by which ER stress is contributing to lung inflammation and damage in the pristane model. ER stress has been shown to contribute to lung inflammation through modulation of the NFκB/HIF-1α pathways (20). ER stress is also known to drive hepatic, endothelial and epithelial cell apoptosis via reactive oxygen-dependent pathways and contribute to disease pathology in various models (34-36). It is important to note that ER stress has also been shown to regulate production of, and sensing of, type I IFNs, resulting in exacerbated IFN-driven responses (37). The IRE1/XBP1 arm of the unfolded protein response has been largely shown to augment type I IFN production via activation of additional stress-responsive proteins such as protein kinase R (PKR), STING and IRF3, or directly by spliced XBP1 binding to the promoter of *Ifnb1* (38-41). In keeping with this, we have found that treatment of alveolar epithelial cells not only drives induction of ER stress markers but also induces the expression of IFN-stimulated genes directly. Thus, although the specific mechanism is currently under investigation, our data suggests that ER stress can directly drive IFNβ production and that NET-induced ER stress may be contributing to IFN induction in the pristane model of SLE.

In summary, the current study shows that ER stress markers are upregulated in the lungs of pristane-treated mice. Targeted deletion of the enzyme PAD4 in myeloid cells reduced lung inflammation and markers of ER stress and IFN-driven responses. Neutrophils isolated from lupus patients and pristane-treated mice could directly upregulate ER stress markers in alveolar epithelial cells and murine lung slices, respectively. Collectively, our data supports role for NETosis and ER stress in driving autoimmune lung inflammation.

## ACKNOWLEDGEMENTS

Authors thank you to funding agencies: Arthritis Foundation AF2017-433570 (Jefferies, PI) and the Leon Fine Translational award, Cedars Sinai Medical Center (Jefferies, PI).

---

**Supplemental table 1:**

Murine primer sequences are listed as follows:

*gapdh*: 5’-CATCAAGAAGGTGGTGAAGC-3’ and 5’-CCTGTTGCTGTAGCTGTATT-3’

*ddit*3: 5’-ACGGAAACAGAGTGGTCAGTGC-3’ and 5’-CAGGAGGTGATGCCCACTGTTC-3’

*hspa*5: 5’-CCTGCGTCGGTGTGTTCAAG-3’ and 5’-AAGGGTCATTCCAAGTGCG-3’

*sxbp1*: 5’-GAGTCCGCAGCAGGTG-3’ and 5’-GTGTCAGAGTCCATGGGA-3’

Total *xbp1*: 5’-GTCCATGGGAAGATGTTCTGG-3’ and 5’-TGGCCGGGTCTGCTGAGTCCG-3’

*ip10*: 5’-GAGTCCGCAGCAGGTG-3’ and 5’-GTGTCAGAGTCCATGGGA-3’

*oas*1: 5’-GCCTGATCCCAGAATCTATGC-3’ and 5’-GAGCAACTCTAGGGCGTACTG-3’

*isg*15: 5’-GGTGTCCGTGACTAACTCCAT-3’ and 5’-CTGTACCACTAGCATCACTGTG-3’

*mx1*: 5’-GATCCGACTTCACTTCCAGATGG-3’ and 5’-CATCTCAGTGGTAGTCAACCC-3’

*ifnb*: 5’-CAGCTCCAAGAAAGGACGAAC-3’ and 5’-GGCAGTGTAACTCTTCTGCAT-3’

*elane*: 5’-GACATGACGAAGTTCCTGGCA-3’ and 5’-CCTTGGCAGACTATCCAGCC-3’

*mpo*: 5’-AGTTGTGCTGAGCTGTATGGA-3’ and 5’-CGGCTGCTTGAAGTAAAACAGG-3’

Human primer sequences are as follows:

*18s*: 5’-GCTTAATTTGACTCAACACGGGA-3’ and 5’-AGCTATCAATCTGTCAATCCTGTC-3’

*ATF4*: 5’-GTTCTCCAGCGACAAGGCTA-3’ and 5’-ATCCTGCTTGCTGTTGTTGG-3’

*CHOP*: 5’-AGAACCAGGAAACGGAAACAGA-3’ and 5’-TCTCCTTCATGCGCTGCTTT-3’

*IFI44L*: 5’-AGCCGTCAGGGATGTACTATAAC-3’ and 5’-AGGGAATCATTTGGCTCTGTAGA-3’

*IL16*: 5’-GCCGAAGACCCTTGGGTTAG-3’ and 5’-GCTGGCATTGGGCTGTAGA-3’

*IL8*: 5’-AAATTTGGGGTGGAAAGGTT-3’ and 5’-TCCTGATTTCTGCAGCTCTGT-3’

*IP10*: 5’-GTGGCATTCAAGGAGTACCTC-3’ and 5’-TGATGGCCTTCGATTCTGGATT-3’

*ISG15*: 5’-ACTCATCTTTGCCAGTACAGGAG-3’ and 5’-CAGCATCTTCACCGTCAGGTC-3’

*RSAD2*: 5’-TTGGACATTCTCGCTATCTCCT-3’ and 5’-AGTGCTTTGATCTGTTCCGTC-3’

*sXBP1*: 5’-CTGAGTCCGAATCAGGTGCAG-3’ and 5’-ATCCATGGGGAGATGTTCTGG-3’

*XBP1*: 5’-TGGCCGGGTCTGCTGAGTCCG-3’ and 5’-ATCCATGGGGAGATGTTCTGG-3’

**Supplemental Figure 1.**
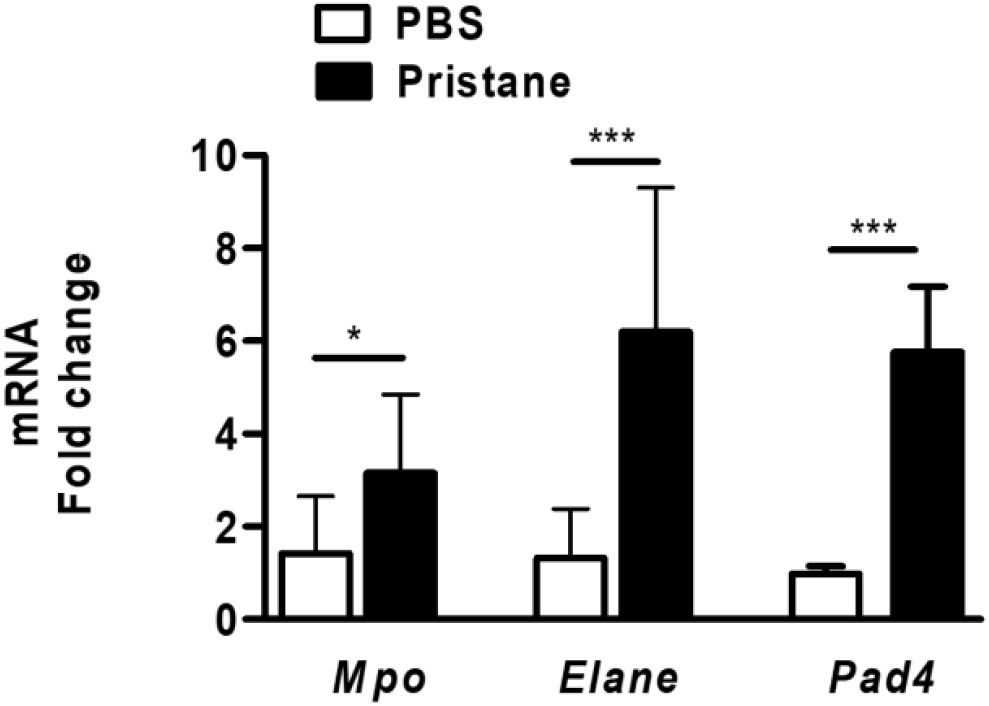
ER stress development coincides with neutrophil infiltration in the lung after prlstane. Expression of neutrophil-associated transcripts were determined by qPCR from whole lungs of WT mice treated with PBS or pristane. Values are the mean ± SD. Statistical significance was determined by unpaired Student’s t-test. \**p* < 0.05,\*\**p* < 0.01, and \*\*\**p ≤* 0.001

**Supplemental Figure 3.**
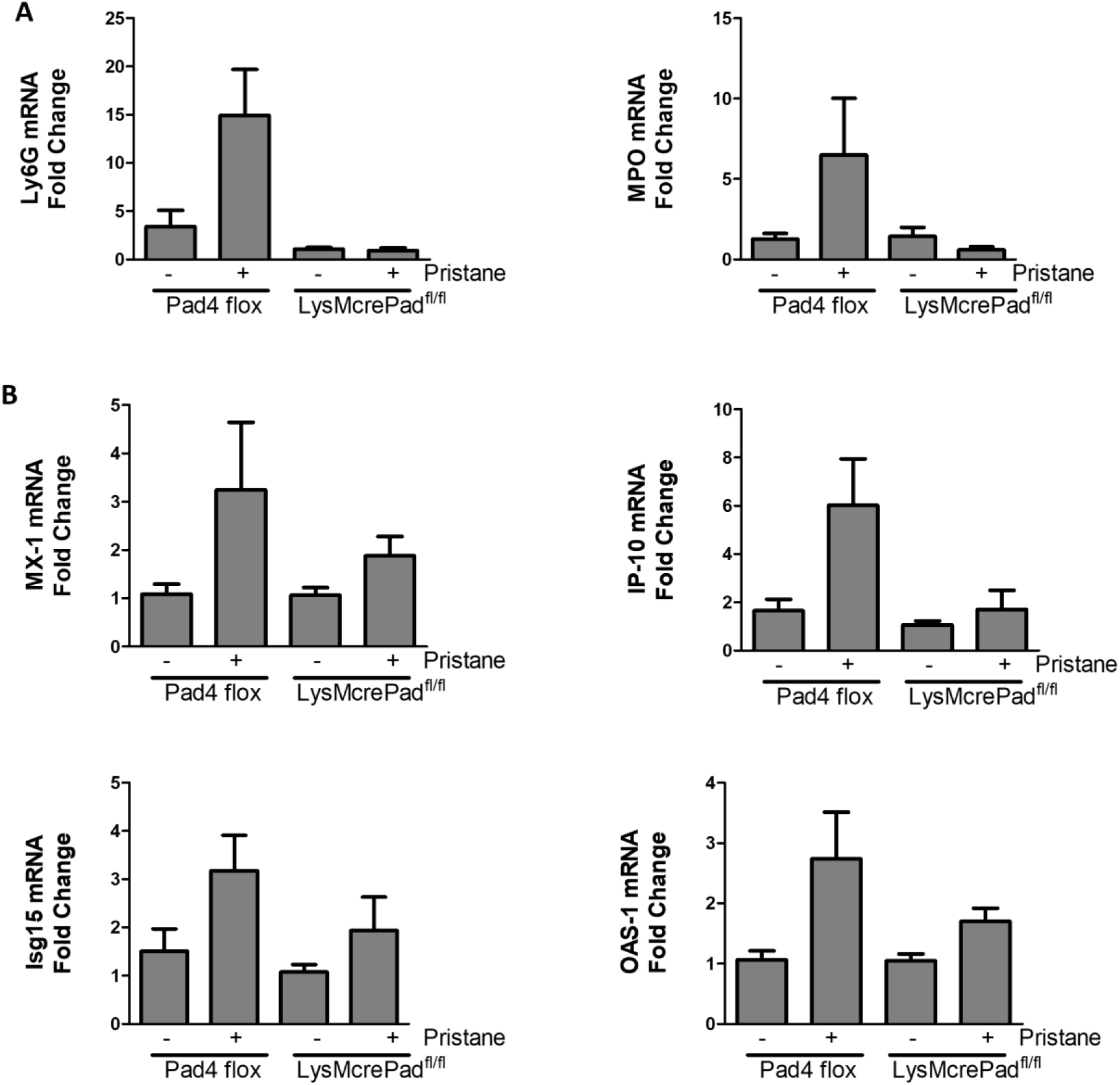
LysMCrePAD4^flfl^ peritoneal cells do not upregulate neutrophil activation genes and are less inflammatory than PAD4^flfl^ cells. Transcripts of **(A)** neutrophil protein markers Ly6G and MPO and (B) ISGs as measured by qPCR of total peritoneal lavage cells 7 days following PBS or pristane injection. values are the mean ± SD. Statistical significance was determined by one-way ANOVA test. \**p* < 0.05,\*\**p* < 0.01, and \*\*\**p* ≤ 0.001

